# Genome-wide CRISPR screens identify noncanonical translation factor eIF2A as an enhancer of SARS-CoV-2 programmed –1 ribosomal frameshifting

**DOI:** 10.1101/2023.01.23.525275

**Authors:** Lian-Huan Wei, Yu Sun, Junjie U. Guo

## Abstract

Many positive-strand RNA viruses, including all known coronaviruses, employ programmed –1 ribosomal frameshifting (–1 PRF) to regulate the translation of polycistronic viral RNAs. However, only a few host factors have been shown to regulate –1 PRF. Through a reporter-based genome-wide CRISPR/Cas9 knockout screen, we identified several host factors that either suppressed or enhanced –1 PRF of SARS-CoV-2. One of these factors is eukaryotic translation initiation factor 2A (eIF2A), which specifically and directly enhanced –1 PRF in vitro and in cells. Consistent with the crucial role of efficient –1 PRF in transcriptase/replicase expression, loss of eIF2A reduced SARS-CoV-2 replication in cells. Transcriptome-wide analysis of eIF2A-interacting RNAs showed that eIF2A primarily interacted with 18S ribosomal RNA near the contacts between the SARS-CoV-2 frameshift-stimulatory element (FSE) and the ribosome. Thus, our results revealed an unexpected role for eIF2A in modulating the translation of specific RNAs independent of its previously described role during initiation.

## INTRODUCTION

A large variety of RNA viruses, including the severe acute respiratory syndrome coronavirus 2 (SARS-CoV-2), contain specific RNA structures that promote programmed –1 ribosomal frameshifting (–1 PRF) and regulate the translation of polycistronic viral RNAs that encode nonstructural proteins (NSPs) essential for viral replication^1^. The 16 NSPs of SARS-CoV-2 are generated from two polyprotein precursors pp1a (NSP1-11) and pp1ab (NSP1-16), which are translated from its largest open reading frame ORF1 (**Figure 1A**) and subsequently cleaved by proteases^2^. Translation of the second half of ORF1 (ORF1b) is preceded by –1 PRF at a heptameric slippery sequence (U UUA AAC) promoted by a downstream three-stem RNA pseudoknot acting as a frameshift-stimulatory element (FSE). The FSE pseudoknot makes specific contacts with the small subunit of the ribosome near the mRNA entrance and causes ribosome stalling^3,4^. Ribosomes that do not frameshift would soon encounter an in-frame stop codon, whereas those that undergo –1 PRF would continue and translate ORF1b, which encodes a variety of essential components of the transcriptase/replicase complex including the RNA-dependent RNA polymerase (RdRp/NSP12).

**Figure 1.**
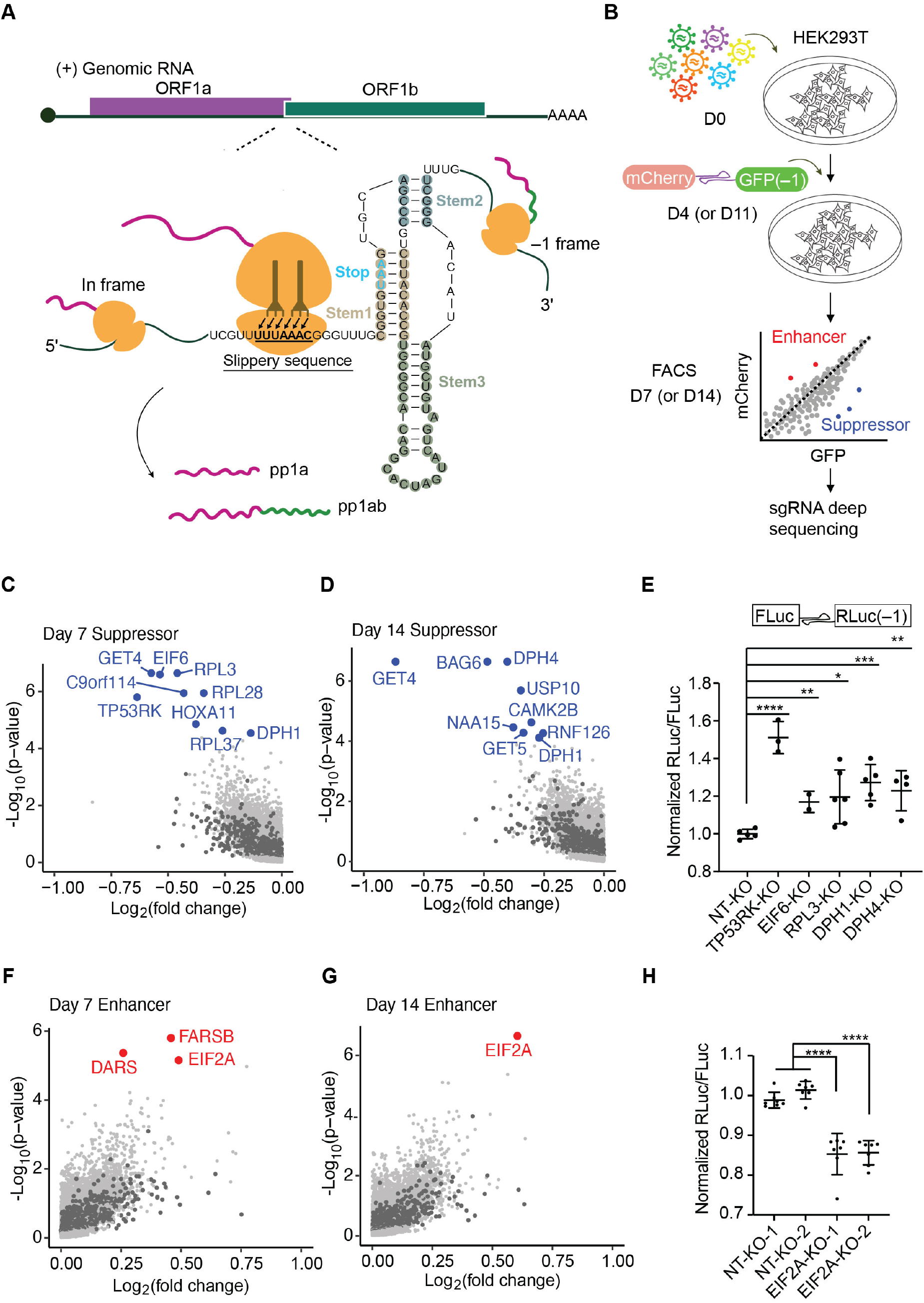
CRISPR/Cas9 knock out screens identify SARS-CoV-2 –1 PRF modulators. **(A**) Schematic illustration of SARS-CoV-2 –1 PRF, showing the locations of ORF1a/1b in the genomic RNA (top) and the key components within FSE (bottom). (**B**) Schematic illustration of the procedure of fluorescent reporter-based genome-wide CRIPSR/Cas9 knock out screen. (**C,D,F,G**) Fold change and statistical significance of modifiers calculated with MAGeCK. Non-targeting negative control sgRNAs are indicated in dark gray. Top-ranking suppressors (**C, D**) and enhancers (**F, G**) are indicated in blue and red, respectively. See also Tables S3, S4. (**E**) Validation of identified –1 PRF suppressors with Fluc-FSE_CoV-2_-Rluc(–1) reporter in HEK293T knockout cell lines. NT, non-targeting sgRNA. (**H**) Validation of EIF2A as a –1 PRF enhancer in HEK293T knockout cell lines. *p < 0.05, **p < 0.01, ***p < 0.001, ****p < 0.0001, two-tailed t test. Data are mean ± SD.

Consistent with the essential role of –1 PRF in viral gene expression, sequence mutations that interfere with –1 PRF can disrupt viral protein stoichiometry and severely compromise viral replication, rendering –1 PRF an attractive target for antiviral development^5–7^. Through an unbiased compound screen, we have recently identified merafloxacin, a fluoroquinolone compound, as a –1 PRF inhibitor that broadly targets betacoronavirus FSEs with a common three-stem pseudoknot fold^7^. We and others have shown that decreasing –1 PRF efficiency by merafloxacin leads to large decreases in virus titer^3,7^, consistent with the notion that SARS-CoV-2 replication strictly requires high –1 PRF efficiency.

In stark contrast to viral RNAs, –1 PRF rarely occurs in cellular mRNAs, with most, if not all, of the few examples originating from retroviruses and retrotransposable elements^8–10^. Indeed, aside from the general mechanisms that ensure reading-frame fidelity during normal translation^11–13^, – 1 PRF suppression by specific factors is part of the host antiviral defense. For example, an interferon-stimulated protein C19orf66/Shiftless inhibits –1 PRF in a wide variety of RNA viruses including human immunodeficiency virus-1 (HIV-1)^14,15^ and SARS-CoV-2^7,16^. Another interferon-stimulated factor zinc-finger antiviral protein (ZAP) has been recently shown to inhibit –1 PRF by directly interacting with SARS-CoV-2 FSE^17,18^. These studies suggest that aside from small molecules, targeting host factors may serve as an alternative strategy for interfering with viral –1 PRF.

In this study, we set out to systematically identify host regulators of –1 PRF through unbiased, genome-wide CRISPR/Cas9 knockout screens, by using our previously developed SARS-CoV-2 –1 PRF reporter. In addition to previously unknown suppressors of –1 PRF, our screen also identified an unexpected role of a noncanonical translation factor eIF2A in promoting –1 PRF, independent of its previously described function during translation initiation.

## RESULTS

### CRISPR screens identified cellular regulators of –1 PRF

We have previously developed and applied a dual fluorescent protein-based reporter to quantify SARS-CoV-2 –1 PRF efficiency^7^. To globally identify cellular factors that modulate –1 PRF of SARS-CoV-2, we first transduced HEK293T cells with lentiviruses expressing Cas9 and an sgRNA library^19^ at a multiplicity of infection (MOI) of <0.3. At each of two subsequent time points (day 4 and 11) after antibiotic selection, cells were transduced with lentiviruses expressing the mCherry-FSE_CoV-2_-GFP (–1) reporter. 3 days after reporter transduction (day 7 and 14), cells with the highest and lowest ~25% GFP/mCherry ratios were harvested by fluorescence-activated cell sorting (FACS), and their sgRNAs were identified and quantified by high-throughput sequencing (**Figure 1B**). As expected, sgRNAs targeting essential genes were more strongly depleted from the pool of cells on day 14 than day 7 (**Figure S1**), indicating the effectiveness of CRISPR/Cas9-mediated knockout (KO) in our screen. While day 7 and day 14 samples yielded largely consistent results (**Figure 1C, D, F, G, Table S1**), more modifiers encoded by essential genes were identified from day 7 samples. For example, sgRNAs targeting essential genes *TP53RK* (p53-regulating kinase), *RPL3* (ribosomal protein L3), and *EIF6* (eukaryotic translation initiation factor 6) significantly enhanced –1 PRF in day 7 samples (**Figure 1C**) before they were depleted in day 14 samples. Their effects on –1 PRF were validated by an orthogonal, dual luciferase-based reporter FLuc-FSE_CoV-2_-Rluc (–1) in HEK293T KO cell lines (**Figure 1E**). Among the overlapping hits between the two time points was *DPH1* (diphthamide biosynthesis protein 1), which encodes the first enzyme in the diphthamide biosynthesis pathway^20,21^. *DPH4* was also identified as a –1 PRF suppressor in day 14 samples. These results were consistent with previous studies showing that the diphthamide modification of translation elongation factor eEF2 suppresses –1 PRF in HIV-1^22,23^. Our screen did not identify ZAP nor Shiftless, presumably because these two interferon-induced genes were not expressed in our unstimulated HEK293T cells. As expected, a majority of the identified –1 PRF modifiers acted as suppressors (**Figure 1C, D, F, G**), with only one candidate enhancer *EIF2A* being consistently identified at both time points. *EIF2A* encodes a noncanonical translation factor eIF2A and should not be confused with the eIF2α subunit (encoded by the *EIF2S1* gene) of the eIF2 complex^24^.

### eIF2A specifically enhanced –1 PRF independent of translation initiation

To validate its effect on –1 PRF, eIF2A was either knocked out or overexpressed in HEK293T cells (**Figure S2A, B**). Consistent with the results from our CRISPR/Cas9 screen, knocking out eIF2A with either of the two sgRNAs significantly decreased –1 PRF efficiency as measured by the dual luciferase reporter (**Figure 1H**). On the contrary, overexpressing eIF2A in HEK293T cells significantly increased –1 PRF but had no effect on luciferases expressed from a construct without an FSE (**Figure 2A**, in-frame control). Similar results were observed when we knocked out or overexpressed eIF2A in Vero E6 cells (**Figure S2C, D**). As expected from the high sequence similarity between SARS-CoV-1 and SARS-CoV-2 FSEs, which differ only by one unpaired nucleotide between Stems 2 and 3, eIF2A overexpression also enhanced –1 PRF of SARS-CoV-1 but not that of HIV-1 (**Figure 2A**). Furthermore, only full-length eIF2A, which contained both an N-terminal WD repeat domain and a C-terminal disordered domain^24^, but neither N-terminal-nor C-terminal-truncated mutants enhanced frameshifting (**Figure 2B**).

**Figure 2.**
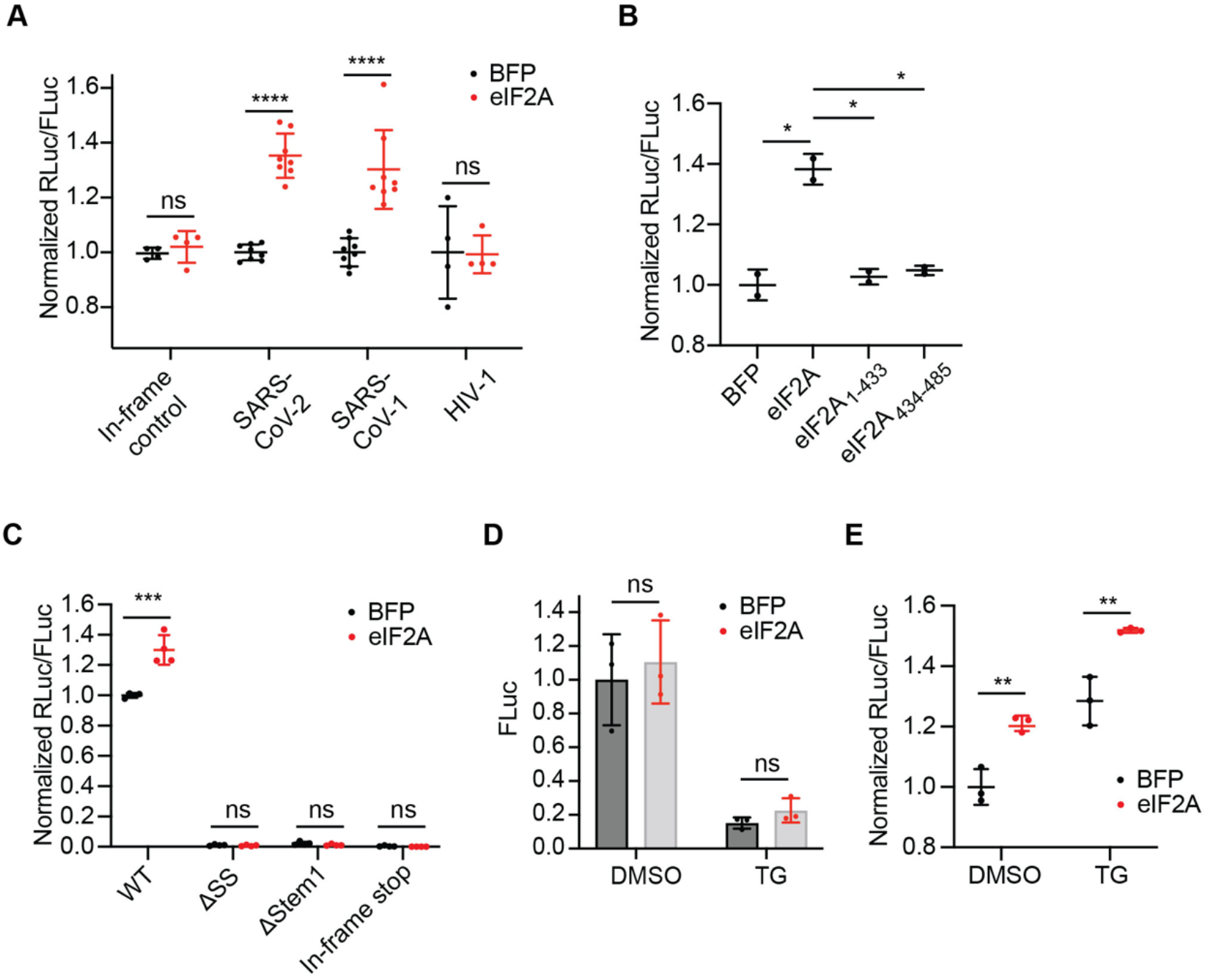
eIF2A enhances SARS-CoV-2 –1 PRF independent of translation initiation. (**A**) Effects of eIF2A overexpression on an in-frame control and –1 PRF reporters for HIV-1, SARS-CoV-1 and SARS-CoV-2 in HEK293T cells. (**B**) Effects of overexpressing eIF2A or its truncated mutants on SARS-CoV-2 –1 PRF in HEK293T cells. (**C**) Effects of eIF2A overexpression on –1 PRF-disrupted control reporters. (**D**) Effects of eIF2A overexpression on overall translation level as measured by FLuc activity in control and thapsigargin (TG)-treated HEK293T cells. (**E**) Effects of eIF2A overexpression on –1 PRF in control and TG-treated HEK293T cells. ns, not significant, *p < 0.05, **p < 0.01, ***p < 0.001, ****p < 0.0001, two-tailed t test. Data are mean ± SD.

The identification of eIF2A as a –1 PRF modifier was surprising because eIF2A, as its name suggested, was not known to regulate translation elongation. Rather, it has been initially identified as an initiator Met-tRNA_i_ delivery factor^25^. A later study, however, has shown that this activity is due to the co-purification of eIF2D and that eIF2A lacks such activity^26^. To investigate the potential confounding effect of eIF2A-dependent changes in translation initiation on –1 PRF, we first tested the possibility of eIF2A acting either through re-initiation in the –1 frame after the in-frame stop codon or through an internal ribosome entry site (IRES), as has been previously discussed^27–30^, thereby increasing RLuc activity. Firstly, disrupting –1 PRF either by deleting the slippery sequence (ΔSS) or by disrupting FSE Stem 1 (ΔStem 1), while leaving the 0-frame stop codon and its immediate flanking sequence intact, completely abolished RLuc activity in both control and eIF2A-overexpressing cells (**Figure 2C; Figure S3**), which ruled out the possibility of re-initiation. Secondly, inserting an additional in-frame stop codon immediately upstream of FSE also abolished the effect of eIF2A (**Figure 2C; Figure S3**), which ruled out the possibility of eIF2A acting through an IRES. These results indicated that the effect of eIF2A strictly required the essential features of –1 PRF and excluded the possibility of unintended initiation in the –1 frame.

To further test the potential influence of eIF2A on the overall translation level of –1 PRF reporter mRNA, we treated cells with an endoplasmic reticulum (ER) stressor thapsigargin (TG), which dramatically decreased global translation including the reporter mRNA^31,32^ (**Figure 2D**). In contrast, eIF2A overexpression did not significantly change the overall translation output from the reporter mRNA in either control or TG-treated cells (**Figure 2D**). While baseline frameshifting efficiency was increased by TG, eIF2A overexpression led to similar increases in frameshifting efficiency in both control and TG-treated cells (**Figure 2E**), suggesting that the effect of eIF2A on –1 PRF could be decoupled from changes in the overall mRNA translation rate. Collectively, our results indicated that the enhancing effect of eIF2A on frameshifting was independent of its previously described role in translation initiation.

### eIF2A directly promoted SARS-CoV-2 –1 PRF in vitro

To determine whether eIF2A directly or indirectly modulated –1 PRF, such as by altering the expression of other cellular factors, we expressed and purified full-length eIF2A protein, as well as its N-terminal or C-terminal truncated variants, from HEK293T cells (**Figure 3A**). We then measured their effects on the frameshifting efficiency of in vitro transcribed reporter mRNAs in rabbit reticulocyte lysate. Consistent with our results from cell-based assays (**Figure 2**), full-length eIF2A specifically enhanced –1 PRF of SARS-CoV-2 in a dose-dependent manner (**Figure 3B**) but had no effect on –1 PRF of HIV-1 (**Figure 3C**). Consistent with their lack of activity in cells, neither N-terminal-nor C-terminal-truncated mutants enhanced –1 PRF in vitro (**Figure 3B-D**). Furthermore, disrupting Stem 1 of the FSE pseudoknot abolished the effect (**Figure 3D**), indicating that eIF2A indeed acted through –1 PRF. These results suggested that eIF2A could directly promote SARS-CoV-2 –1 PRF without changing the expression of other cellular factors.

**Figure 3.**
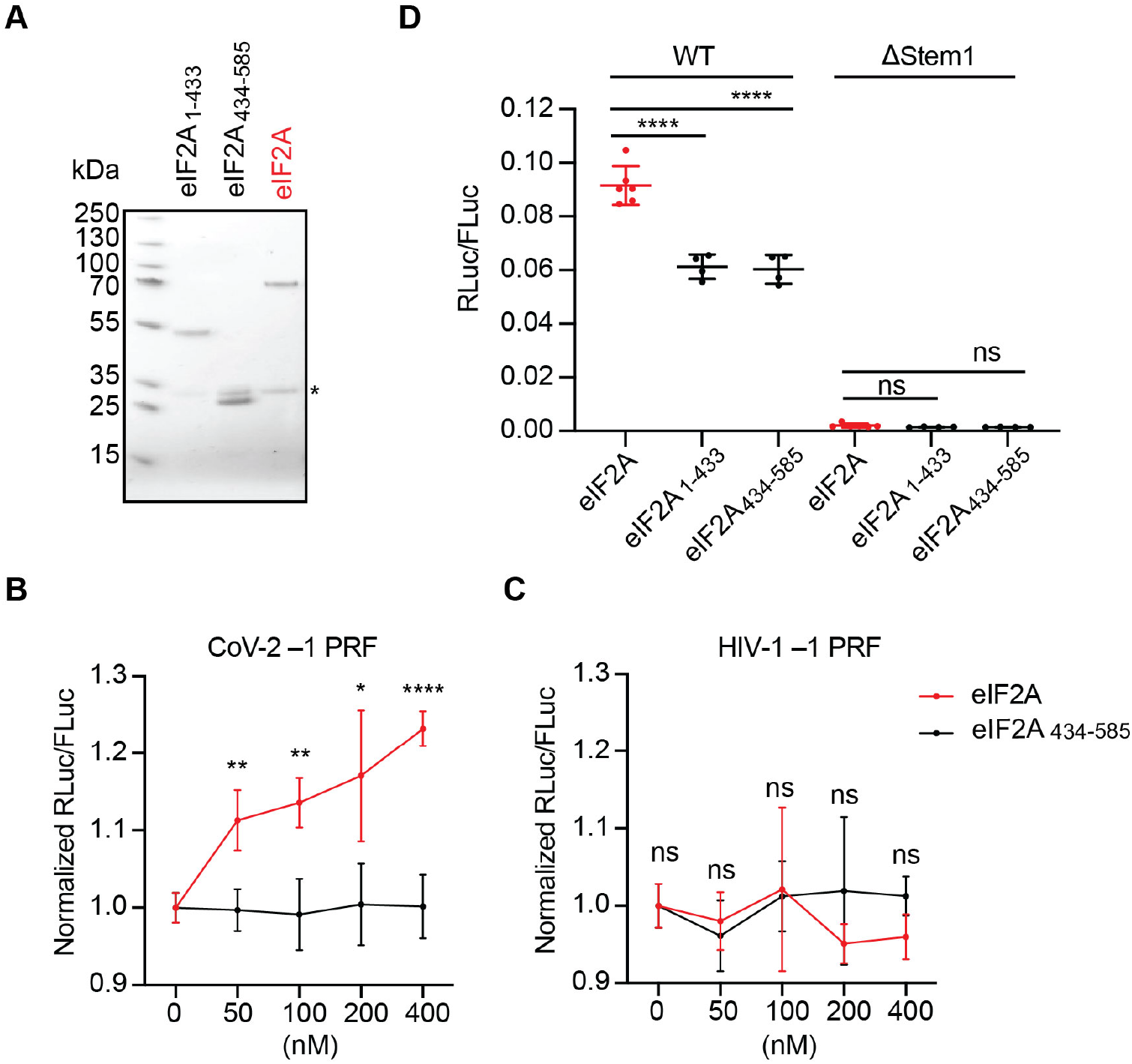
eIF2A directly promotes SARS-CoV-2 –1 PRF. (**A**) Commassie blue-stained SDS-PAGE showing recombinant full-length eIF2A (65 kDa), a C-terminal-truncated EIF2A_1-433_ (48 kDa), and a N-terminal-truncated EIF2A_434-585_ (17 kDa). *, Tobacco Etch Virus (TEV) protease (28 kDa). (**B, C**) Dose-dependent effects of recombinant eIF2A and mutant on SARS-CoV-2 (**B**) and HIV-1 (**C**) –1 PRF in rabbit reticulocyte lysate. Data were from four biological replicates and normalized to BSA. (**D**) Effects of recombinant eIF2A or mutants on WT and ΔStem 1 reporters in rabbit reticulocyte lysate. ns, not significant, *p < 0.05, **p < 0.01, ****p < 0.0001, two-tailed t test. Data are mean ± SD.

### Loss of eIF2A reduced SARS-CoV-2 replication

By using quantitative proteomics, a previous study has shown that eIF2A protein abundance is increased in SARS-CoV-1 and SARS-CoV-2 infected A549 cells compared to mock-treated cells, while its mRNA level remains unchanged^33^ (**Figure S4A**). Another proteomics study using Caco-2 cells has also shown that eIF2A protein abundance is progressively increased after SARS-CoV-2 infection^34^ (**Figure S4B**). A third study by profiling ribosome occupancy has shown an increase in eIF2A mRNA translational efficiency after SARS-CoV-2 infection (**Figure S4C**)^35^. Consistent with these studies, we found that eIF2A protein abundance was increased in SARS-CoV-2-infected Vero E6 cells (**Figure 4A, B**), while its mRNA level remained unchanged (**Figure 4C**). These results, together with the effect of eIF2A on –1 PRF, prompted us to examine the role of endogenous eIF2A in SARS-CoV-2 replication. Consistent with the crucial role of –1 PRF on SARS-CoV-2 transcriptase/replicase expression, the intracellular abundance of subgenomic (N) and genomic (ORF1a) RNAs were both decreased in EIF2A KO compared to control Vero E6 cells (**Figure 4D**). Plaque assays showed a similar decrease in the yield of infectious viral particles (**Figure 4E**). To confirm the role of eIF2A in promoting –1 PRF, we measured the protein abundance of NSP8 (encoded in ORF1a) and NSP12/RdRp (encoded in ORF1b). While both NSP8 and NSP12 protein levels were decreased in EIF2A KO cells as expected from the overall reduced viral titer, NSP12 showed a larger reduction than NSP8 in EIF2A KO cells (**Figure 4F**), consistent with a decrease in –1 PRF efficiency. Together, these results indicated that decreasing –1 PRF efficiency by the loss of eIF2A reduced SARS-CoV-2 replication, further supporting the previous findings by us and others showing that coronavirus replication strictly depends on optimal –1 PRF efficiency^3,5,7^.

**Figure 4.**
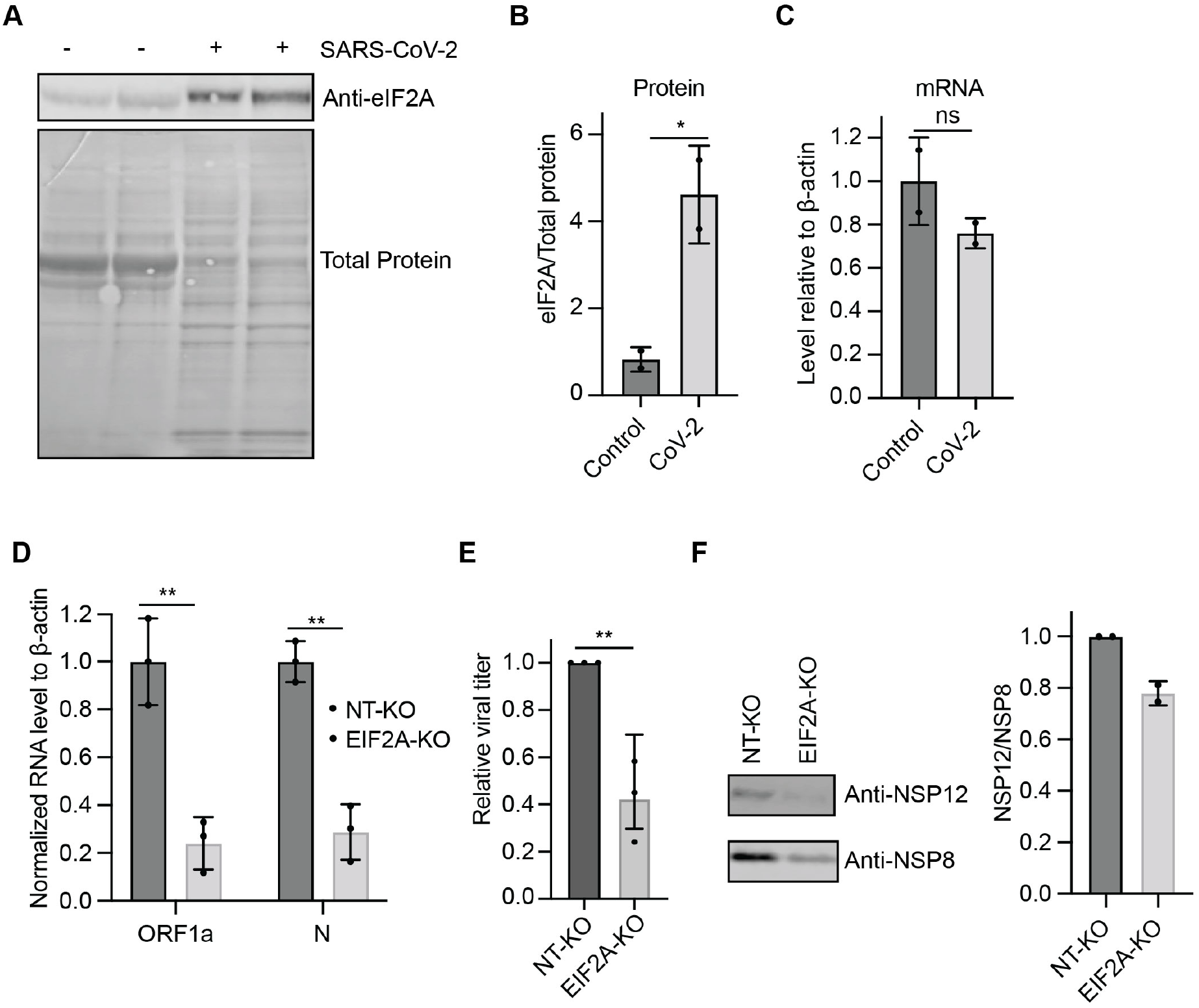
Loss of eIF2A reduces SARS-CoV-2 viral replication. (**A**) Western blots showing the abundance of eIF2A in control and SARS-CoV-2-infected Vero E6 cells. Total protein was stained with Ponceau S solution. (**B**) Quantification of eIF2A protein abundance normalized to total protein abundance. (**C**) Relative abundance of eIF2A mRNA in control and infected Vero E6 cells. (**D**) Relative abundance of ORF1a (genomic RNA) and N (genomic and subgenomic RNA) in control and EIF2A-KO Vero E6 cells 48 hours after infection. (**E**) SARS-CoV-2 viral yield from control and EIF2A-KO Vero E6 cells 48 hours after infection. (**F**) Western blots (*left*) and quantification (*right*) showing the abundance of NSP8 and NSP12 in control and EIF2A-KO Vero E6 cells 48 hours after infection. ns, not significant, *p < 0.05, **p < 0.01, two-tailed t test. Data are mean ± SD.

### eIF2A interacted with 18S rRNA near FSE interactions

To understand the mechanism of eIF2A promoting –1 PRF, we performed enhanced crosslinking and immunoprecipitation (eCLIP)^36^ followed by high-throughput sequencing to identify the RNA interactome of eIF2A in HEK293T cells. We first assessed the previously described role of eIF2A as an alternative delivery factor for initiator Met-tRNA_i_ and Leu-tRNA-CAG^25,26,37,38^. Although various modifications in tRNAs impeded full-length cDNA synthesis, we still obtained a small fraction (1-2%) of reads from both input and eIF2A IP samples uniquely mapped to tRNAs (**Figure 5A, Table S2**). Met-tRNA_i_ and Leu-tRNA-CAG accounted for only ~0.07 % and ~0.02% of all uniquely mapped eIF2A eCLIP reads or ~3% or ~1% of all tRNA-mapping reads, respectively. Importantly, similar to GFP controls (**Figure S5**), the composition of eIF2A-bound tRNAs, including Met-tRNA_i_ and Leu-tRNA-CAG, largely reflected that of the input prior to eIF2A IP (**Figure 5B**), suggesting that most tRNA reads represented nonspecific binding.

**Figure 5.**
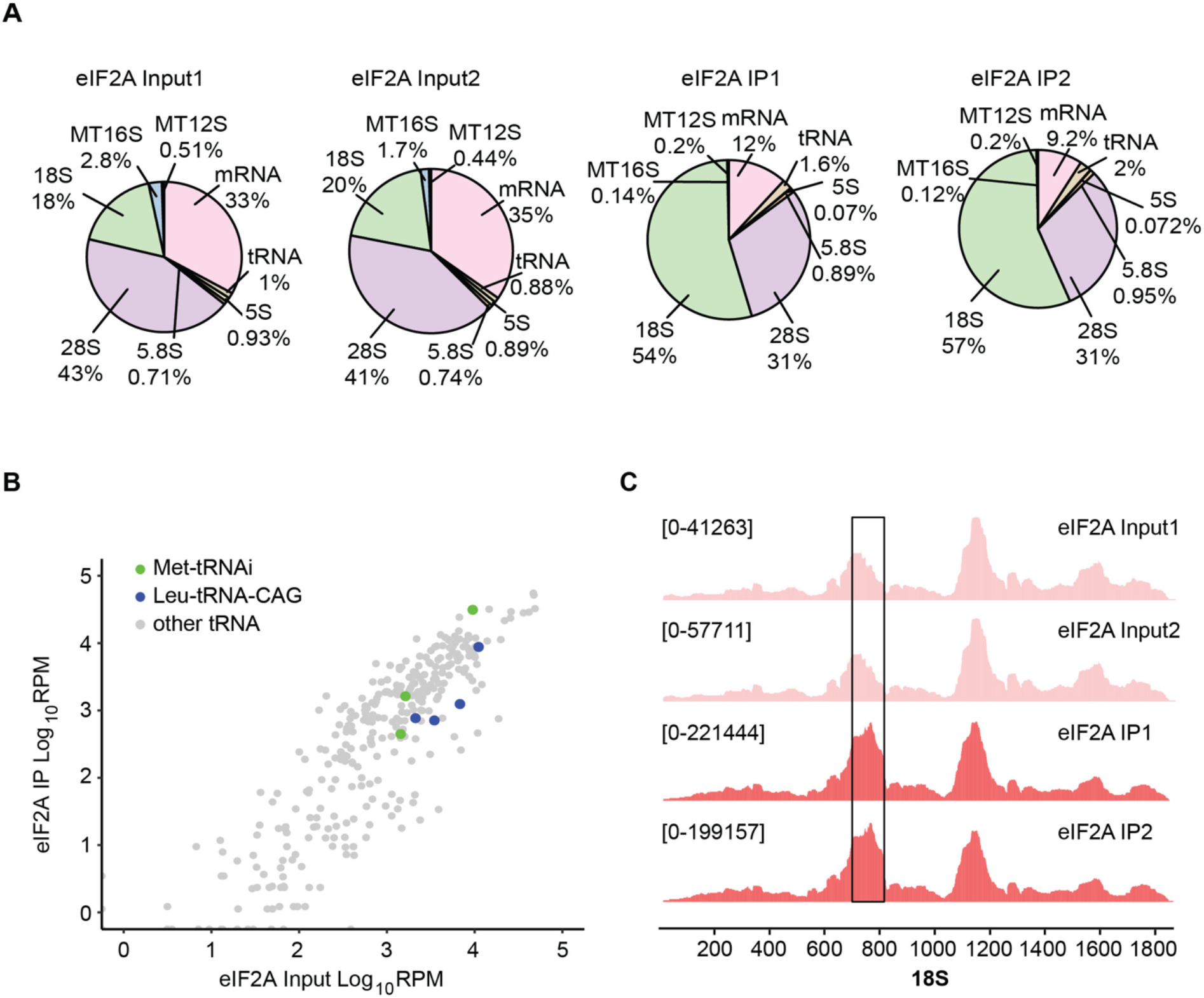
eIF2A specifically binds 18S rRNA near FSE interactions. (**A**) Biotypes of eCLIP reads from eIF2A input and IP samples. Percentage of each biotype is shown. (**B**) Read coverage on 18S rRNA. A putative binding site of eIF2A is indicated by a box. See results of other rRNAs in Figure S3. (**C**) tRNA read counts of eIF2A input and IP samples. Met-tRNA_i_ and Leu-tRNA-CAG species are indicated in green and blue, respectively.

In contrast to tRNAs, mRNAs, and other cytoplasmic (5S, 5.8S, and 28S) and mitochondrial (12S and 16S) rRNAs (**Figure 5A; Figure S6**), 18S rRNA was strongly enriched after eIF2A IP, with a majority (54-57%) of eCLIP reads mapped to 18S rRNA, compared to 18-20% from the input RNAs (**Figure 5A**). While high read coverage was observed across the entire 18S rRNA, eIF2A bound more strongly to a region between helices 21es6B and 21es6C of the central domain^39^ (**Figure 5C; Figure S7**). This region is spatially adjacent to the mRNA entrance and the specific contacts between the SARS-CoV-2 FSE pseudoknot and 18S rRNA^3^ (**Figure S7A, B**), suggesting that eIF2A may promote –1 PRF efficiency in part through its direct rRNA binding affecting the specific interactions between SARS-CoV-2 RNA and the ribosome.

## DISCUSSION

Motivated by recent studies identifying two host factors suppressing –1 PRF in RNA viruses including SARS-CoV-2 by using a candidate-based approach^14,15,17,18^, we set out to comprehensively identify such factors through genome-wide CRISPR/Cas9-based genetic screens. We took advantage of our recently developed and well characterized fluorescent protein-based and luciferase-based SARS-CoV-2 –1 PRF reporters, which allowed us to quantify –1 PRF efficiency in single cells and in bulk, respectively^7^. Our screens were performed in unstimulated HEK293T cells, which may explain why we did not identify the two known suppressors Shiftless and ZAP, both encoded by interferon-stimulated genes. Additional false negatives may include other cell-type-specific factors that are not expressed in HEK293T cells as well as genes that are strictly essential for cell survival and therefore their sgRNAs would drop out even at the earlier time point (7 days after Cas9/sgRNA expression). Nevertheless, our screens identified novel –1 PRF suppressors including some that caused survival defects when knocked out. Among our identified suppressors are diphthamide synthesis enzymes DPH1 and DPH4 (**Figure 1C-E**), which modify elongation factor eEF2 and have previously been shown to suppress –1 PRF of HIV-1^22,23^, suggesting that loss of diphthamide modification on eEF2 generally compromised translation fidelity. Similarly, loss of essential ribosome protein RPL3 also increased –1 PRF (**Figure 1C, E**), whereas the mechanisms of other newly identified –1 PRF suppressors are currently unknown.

A previous study has shown that the encephalomyocarditis virus (EMCV) 2A protein, which inhibits cap-dependent translational initiation in the host, directly binds its FSE and enhances –1 PRF^40^. However, very few cellular mRNAs undergo –1 PRF, and no cellular factors have been shown to enhance –1 PRF. To our surprise, our screens at two separate time points yielded consistent results showing that knocking out eIF2A decreased –1 PRF, suggesting that eIF2A might act as a –1 PRF enhancer (**Figure 1F, G**). To ensure that this phenotype was not due to its previously described role in translation initiation, we showed that (1) the effect of eIF2A required the slippery sequence, the pseudoknot, and in-frame translation; (2) eIF2A overexpression did not change overall translation rate of the –1 PRF reporter mRNA; and (3) globally decreased translation initiation by ER stress did not change the effect of eIF2A on –1 PRF (**Figure 2C-E**). These results indicated that the effect of eIF2A on –1 PRF was independent of any changes in translation initiation.

Unlike Shiftless and DPH enzymes, which broadly inhibit –1 PRF, eIF2A promoted –1 PRF of SARS-CoV-1 and −2 but not that of HIV-1 both in vitro and in cells (**Figure 2A; Figure 3C**), arguing against a general role of eIF2A in translation fidelity. In addition, FSE-specific enhancement of –1 PRF by eIF2A did not require changes in the expression of other cellular factors, as recombinant eIF2A could promote –1 PRF in an in vitro translation system (**Figure 3**). Consistent with a direct effect of eIF2A on –1 PRF, eCLIP analysis showed that eIF2A interacted with 18S rRNA in the highly variable V4 region of its central domain near the mRNA entrance (**Figure 5A, C; Figure S6; Figure S7**), suggesting that it may form specific contacts with both the ribosome and the FSE pseudoknot. Future studies on the ribosome-eIF2A-mRNA complex should reveal the potential structural changes that promote –1 PRF.

Recent studies have elucidated several mechanisms by which SARS-CoV-2 stifles the innate immune response in host cells, including shutting down host-cell gene expression by inhibiting the processing, nuclear export, and translation of cellular mRNAs^40–42^. Surprisingly, we and others observed a large increase in eIF2A protein abundance in SARS-CoV-2-infected cells^33,34^, while its mRNA level remained unchanged, suggesting either that eIF2A mRNA was more efficiently translated or that eIF2A protein became more stable upon infection. Interestingly, a recent study has shown a time-dependent increase in eIF2A mRNA translational efficiency after SARS-CoV-2 infection^35^ (**Figure S4C**). The same study has also shown that –1 PRF efficiency, as measured by the relative ribosome footprint densities between ORF1A and ORF1B, is significantly increased at later time points during infection. Collectively, these results raised an intriguing possibility that eIF2A may play a role in the temporal regulation of SARS-CoV-2 gene expression.

The physiological role of eIF2A has been debated since its initial identification as an initiator Met-tRNA_i_ delivery factor^25,26^. Subsequent studies have proposed additional roles of eIF2A in promoting translation initiation at noncanonical start codons such as CUG^37,43^, in part by delivering Leu-tRNA-CAG to the 40S ribosome^37^. Consistent with a lack of eIF2A KO effect on cell growth in our screen, eIF2A KO is well tolerated during mouse development and only causes sex-specific metabolic changes later in life^44,45^, indicating that eIF2A is not required for general translation. Our eIF2A eCLIP results showed roughly background-level binding of both Met-tRNA_i_ and Leu-tRNA-CAG (**Figure 5B; Figure S5**). This may be either due to the low binding affinity of eIF2A for Met-tRNA_i_^26^ or the binding being strictly dependent on the presence of an mRNA template as previously suggested^38,46^, resulting in a surplus of tRNA-free eIF2A. Instead, we found that eIF2A mostly bound 18S rRNA, which accounted for the majority of eIF2A eCLIP reads (**Figure 5A**), with a particular enrichment near the mRNA entrance (**Figure 5C, Figure S7**). Based on these results, our findings on the effects of eIF2A on –1 PRF may reflect its physiological function in regulating translation in an mRNA-specific manner.

## METHODS

### Reporter lentivirus preparation

Lipofectamine 2000 (Life Technologies) was used for transfection according to the manufacturer’s instructions. For each T175 flask, 15 μg mCherry-FSE_CoV-2_-GFP (–1) reporter^7^, 15 μg pMD.2G, and 15 μg psPA2 were mixed in 4 mL OptiMEM (Life Technologies). After 5 mins, 100 μL of Lipofectamine 2000, diluted in 4 mL OptiMEM, was added to the plasmid mixture. The mixture was incubated at room temperature for 20 mins before being added to 60-70% confluent HEK293T cells. The medium was replaced by fresh medium after 4-6 hours. After 48 hours, viruses were collected and concentrated using Lenti-X Concentrator (Takara, 631232).

### CRISPR knockout screen

The Human GeCKOv2 CRISPR knockout pooled library^19^ (Addgene 1000000048), which contains 6 sgRNAs for each gene and 2,000 non-targeting control sgRNAs, was used in our screen, following the previously described procedure^47^ with minor modifications. For each of the two sub-libraries A and B, approximately 150 million HEK293T cells were transduced at MOI < 0.3 in the presence of 8 μg/mL polybrene on day 0. On day 1, the transduced cells were seeded into medium containing 0.5 μg/mL puromycin. Cells were infected with the mCherry-FSE_CoV-2_-GFP(–1) reporter lentivirus on day 4 (or day 11). Approximately 10 million cells from the top and bottom 25% GFP/mCherry ratios were collected using FACS on day 7 (or day 14). Genomic DNA was extracted using the Zymo Research Quick-gDNA Midiprep Plus kit (D4075). sgRNA sequences were amplified by PCR and subjected to Illumina NovaSeq sequencing. Approximately 20 million reads were collected for each sample. Gene-level enrichment was calculated and ranked using MAGeCK^48^ (**Table S1**).

### Protein purification

EIF2A cDNA was cloned from pDONR223_EIF2A_WT (Addgene, 82111). EIF2A-TEV-GFP, EIF2A_1-433_-TEV-GFP, and EIF2A_434-585_-TEV-GFP were expressed in HEK293T cells. Cells from two 15-cm dishes were pelleted and resuspended in 800 μL of ice-cold lysis buffer (10 mM Tris-HCl, 150 mM NaCl, 0.5 mM EDTA, 0.5% NP-40, pH 7.5) containing 1 mM DTT, 1× protease inhibitor (Millipore, 539134), and 1 mM PMSF. Samples were rotated at 4 °C for 30 mins. After centrifuging at 15,000 rpm for 10 mins at 4 °C, the supernatant was added to 1.2 mL ice-cold dilution buffer (10 mM Tris-HCl, 150 mM NaCl, 0.5 mM EDTA, pH 7.5) containing 1 mM DTT, 1× protease inhibitor, and 1 mM PMSF. 100 μL GFP-Trap magnetic agarose beads (Chromotek) were washed twice with 500 μL ice-cold dilution buffer. The diluted sample was added to the equilibrated beads and rotated end-over-end for 2 hours at 4°C. The beads were then washed with 1 mL high-salt wash buffer (10 mM Tris-HCl, 500 mM NaCl, 0.5 mM EDTA, 0.05% NP-40, pH 7.5) three times, 1 mL wash buffer (10 mM Tris-HCl, 150 mM NaCl, 0.5 mM EDTA, 0.05% NP-40, pH 7.5) three times, and then resuspended in 592 μL of 1× AcTEV cleavage buffer (50 mM Tris-HCl, 0.5 mM EDTA, pH 8.0) containing 1 mM DTT. 8 μL AcTEV protease (Invitrogen, 12575015) was added to the sample and incubated at 30°C for 1 hour with shaking at 1000 rpm. After TEV cleavage, the supernatant was collected, and the beads were further washed twice with 500 μL wash buffer and twice with 500 μL dilution buffer. All eluted fractions were combined and concentrated using centrifugal filter (Millipore) in storage buffer (10 mM Tris-HCl, pH 8.0, 150 mM NaCl, 1 mM DTT, 5% glycerol).

### In vitro translation

In vitro translation was performed using a rabbit reticulocyte lysate system (Promega, L4960). 1.5 μL of purified eIF2A protein or BSA was added to 10 μL rabbit reticulocyte lysate reaction mix, followed by the addition of 1 μL of 2 nM in vitro transcribed luciferase reporter mRNA, prepared using the HiScribe T7 ARCA mRNA Kit (NEB, E2060). The translation reactions were incubated at 30°C for 40 mins. Luciferase activities were measured by Dual-Glo luciferase assays (Promega) and normalized to BSA.

### Cellular –1 PRF assays

To measure the frameshift efficiency in cells, Fluc-FSE_CoV-2_-Rluc (–1) reporter plasmid DNAs were transfected using Lipofectamine 2000 (Thermo Fisher, 11668030). After 48 hours, luciferase activities were measured using the Dual-Glo Luciferase Assay System (Promega, E2920) and the frameshift efficiency was calculated as the ratio of RLuc to FLuc.

### SARS-CoV-2 infection

SARS-CoV-2 infection was performed as previously described^7^. Briefly, cells were seeded at 10^5^ cells/well in 6-well plates. After 24 hours, the cells were incubated with viral stock at MOI of 0.002 for 1 hour, and then washed twice with PBS. 2 days after infection, both infected cells and the media were collected and stored at –80°C. Viral genomic RNA and sub-genomic RNA were extracted from infected cells using TRIzol (Invitrogen) and quantified with RT-qPCR. Plaque assays were performed as previously described^7^.

### RT-qPCR

Total RNA was extracted from cells using TRIzol (Invitrogen, 15596026) and treated with TURBO DNase (Thermo Fisher, AM2238). RT-qPCR was performed using Luna Universal One-Step RT-qPCR reagents (NEB, E3005) and the primers listed in **Table S3**.

### eCLIP-seq

A 10-cm plate of HEK293T cells at 60-70% confluency was co-transfected with the mCherry-FSE_CoV-2_-GFP(–1) reporter and GFP or EIF2A-GFP plasmids. After 48 hours, cells were washed with 10 mL ice-cold 1× DPBS. Cells in 2 mL cold 1× DPBS were crosslinked using 254 nm UV at 400 mJ/cm^2^ on ice. After cell lysis, eCLIP libraries were constructed using the RBP-eCLIP kit (ECLIPSEbio) according to the manufacturer’s instructions. The libraries were sequenced by Illumina NovaSeq and 30 million reads were collected for each sample. eCLIP data processing was performed as previously described^36^ with modifications. Reads were demultiplexed and unique molecular identifiers were extracted using UMI tools^49^. Adapters were trimmed using Cutadapt 3.2^50^, and only reads longer than 18 nt were kept. Trimmed reads were first aligned to non-redundant rRNA and tRNA references^51,52^ with STAR 2.7^53^. Reads that did not map to rRNA and tRNA were then aligned to mRNA references with STAR. After alignment, reads were deduplicated using UMI tools.

## ACKNOWLEDGEMENTS

We thank Ivan Lomakin, Swapnil Chandrakant Devarkar, Yong Xiong, Paulina Pawlica, Sarah Slavoff, and all members of the Guo lab for helpful discussions. J.U.G. is a Klingenstein-Simons Fellow in Neuroscience and a New York Stem Cell Foundation–Robertson Investigator.

## Supplementary Materials

**Table S1** MAGeCK statistics of CRISPR screens.

**Table S2** eIF2A eCLIP results for tRNAs

**Table S3.**
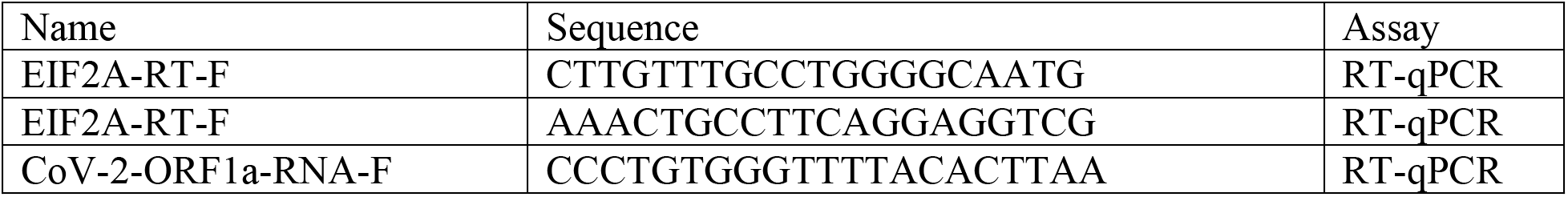
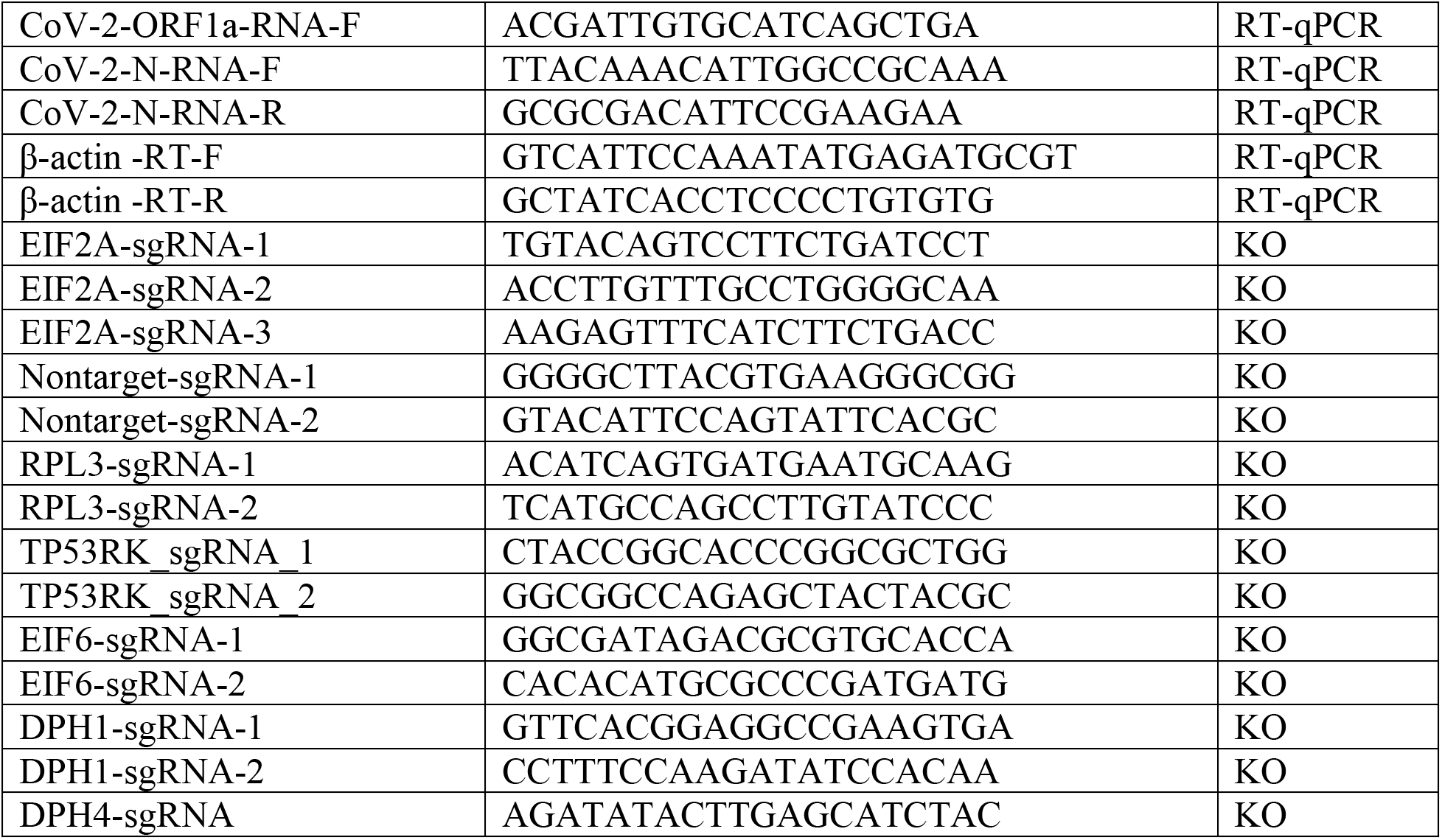
Oligonucleotide sequences used in this study

**Table S4.**
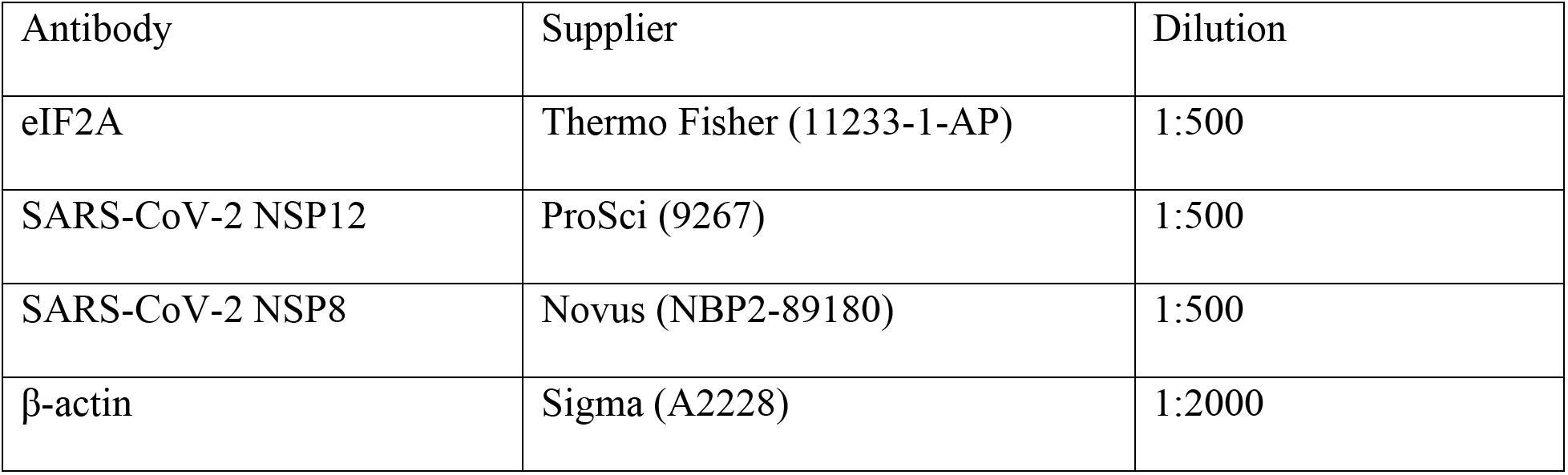
Antibodies used in this study.

| Antibody | Supplier | Dilution |
| --- | --- | --- |
| eIF2A | Thermo Fisher (11233-1-AP) | 1:500 |
| SARS-CoV-2 NSP12 | ProSci (9267) | 1:500 |
| SARS-CoV-2 NSP8 | Novus (NBP2-89180) | 1:500 |
| β-actin | Sigma (A2228) | 1:2000 |

**Figure S1.**
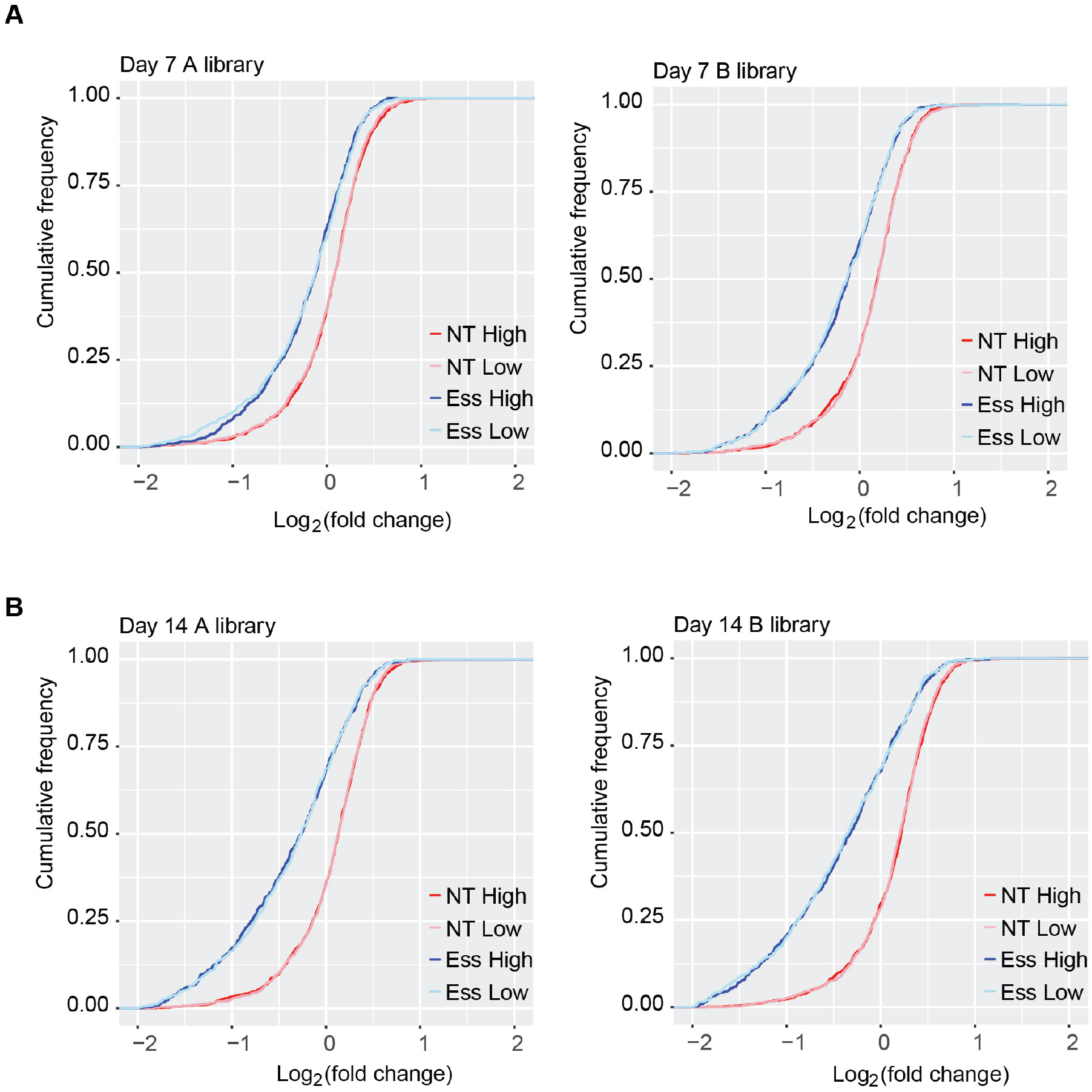
Gradual depletion of sgRNAs for essential genes from the population. Cumulative distribution of changes in the abundance of non-targeting and essential gene^54^ sgRNAs on day 7 (**A**) and day 14 (**B**) samples relative to the sgRNA library prior to transfection. NT High, non-targeting sgRNAs in GFP-to-mCherry-high sample; NT Low, non-targeting sgRNAs in GFP-to-mCherry-low sample; Ess High, sgRNAs for essential genes in GFP-to-mCherry-high sample; Ess Low, sgRNAs for essential genes in GFP-to-mCherry-low sample.

**Figure S2.**
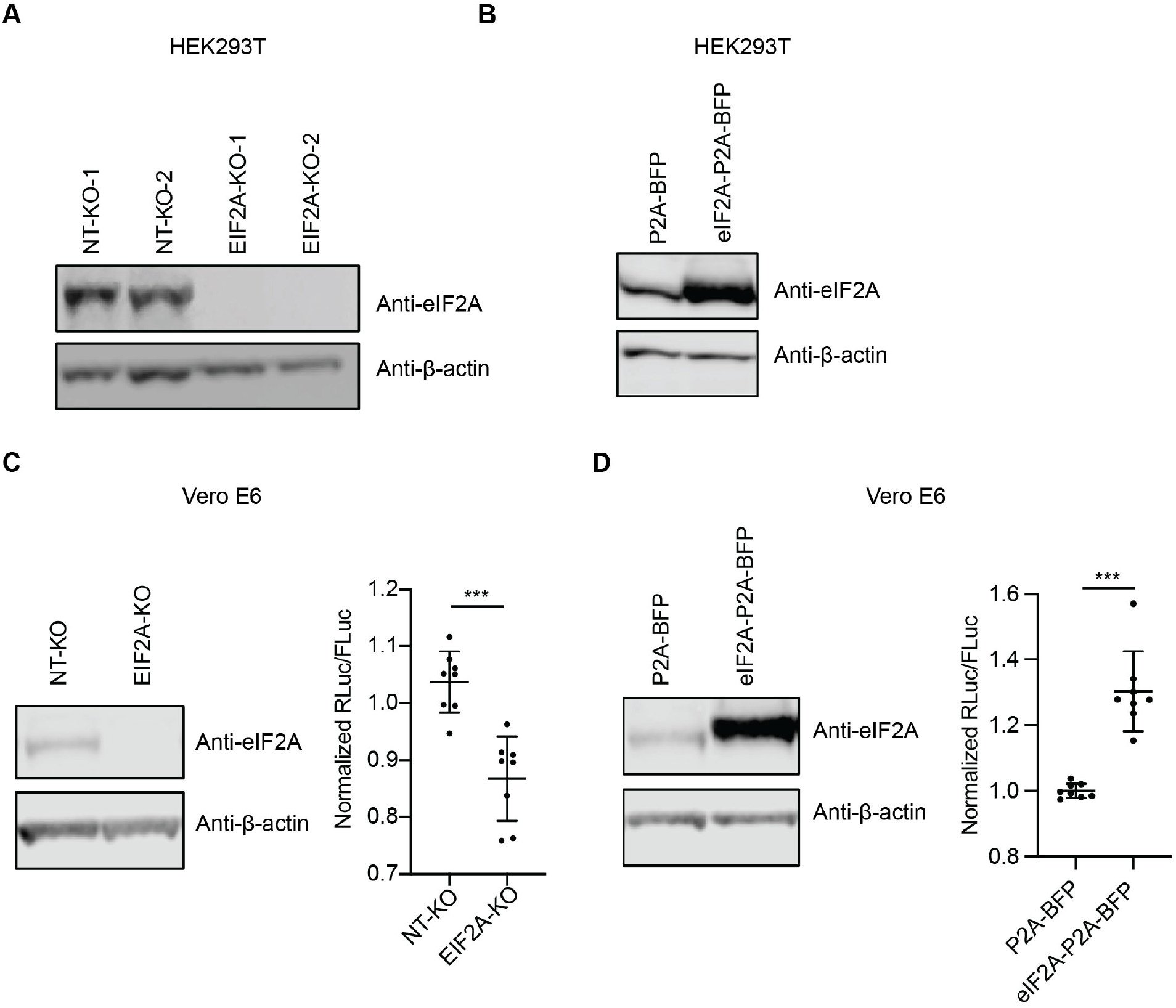
Additional validation of eIF2A as a –1 PRF enhancer. (**A**) Western blots showing the abundance of eIF2A and β-actin in control and EIF2A-KO HEK293T cell lines. (**B**) Western blots showing the abundance of eIF2A and β-actin in control and eIF2A-overexpressing HEK293T cells. **(C) Left:** Western blots showing the abundance of eIF2A and β-actin in control and EIF2A-KO Vero E6 cell lines. **Right:** Effect of eIF2A knockout on –1 PRF in Vero E6 cells. (**D**) **Left:** Western blots showing the abundance of eIF2A and β-actin in control and eIF2A-overexpressing Vero E6 cells. **Right:** Effect of eIF2A overexpression on –1 PRF in Vero E6 cells. ***p < 0.001, two-tailed t test. Data are mean ± SD.

**Figure S3.**
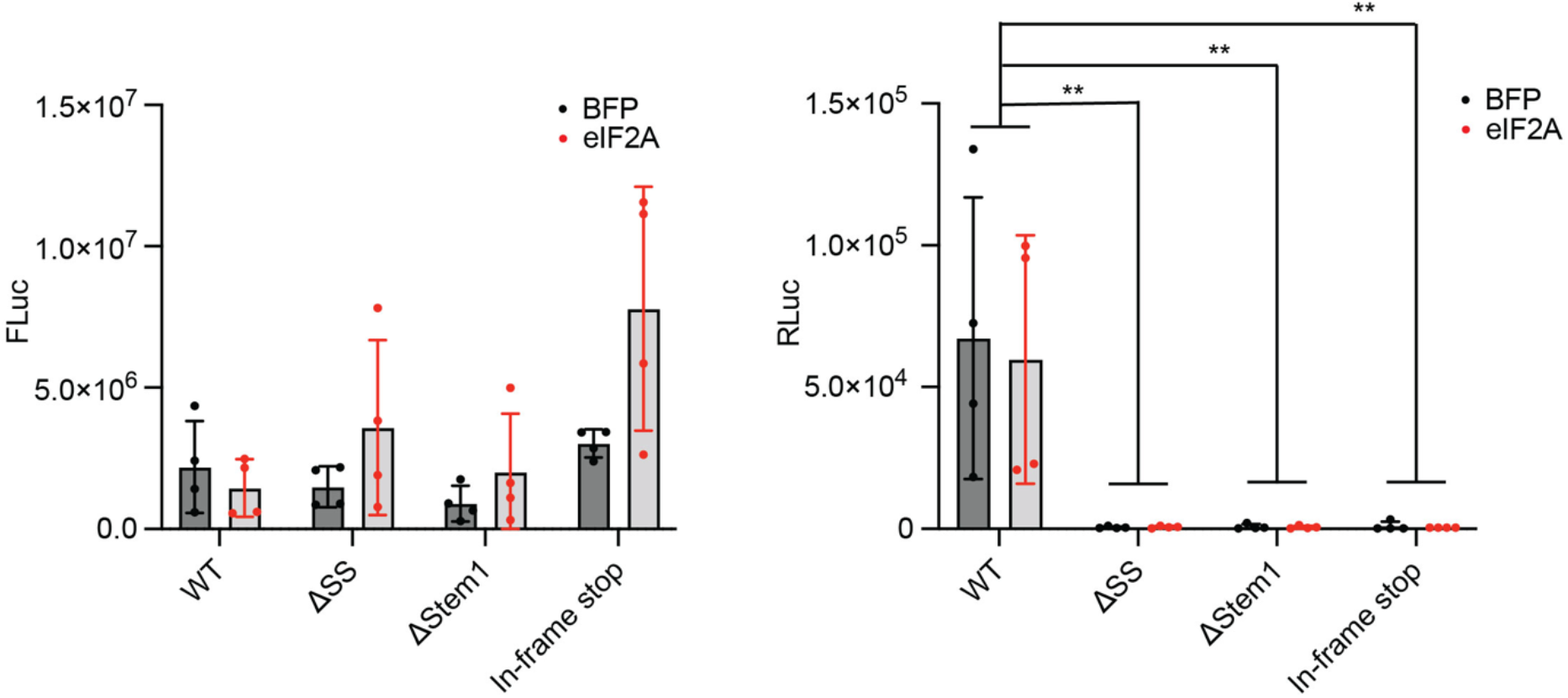
Effects of eIF2A overexpression on FLuc and Rluc activity in –1 PRF-disrupted control reporters.

**Figure S4.**
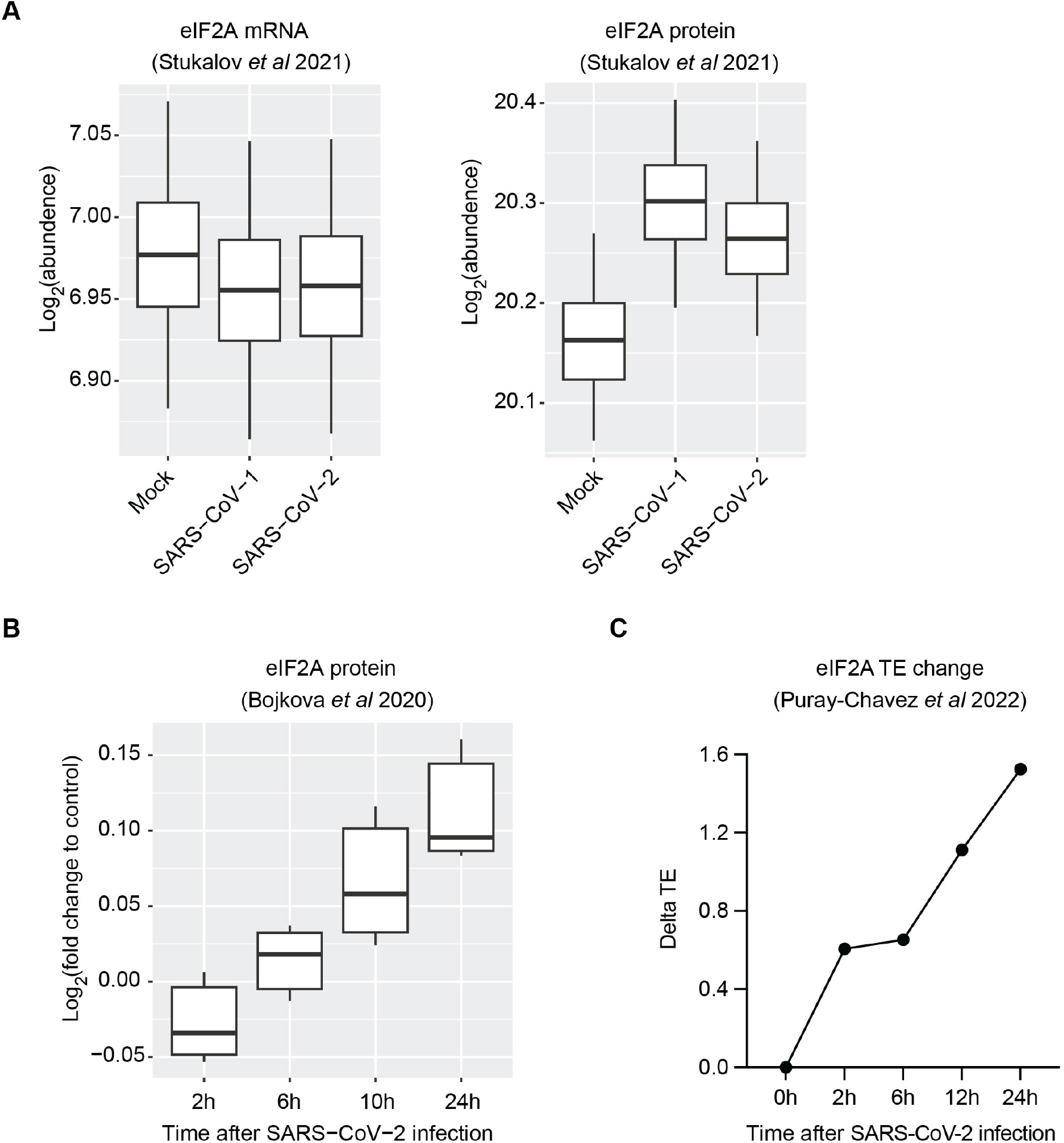
eIF2A mRNA, protein levels, and translational efficiency changes upon viral infection. (**A-B**) Boxplots reproduced using datasets from Stukalov et al. (2021)^33^ (**A**) and Bojkova et al. (2020)^34^ (**B**). (**C**) Changes in the translational efficiency (TE) of eIF2A mRNA using data from Puray-Chavez et al. (2022)^35^.

**Figure S5.**
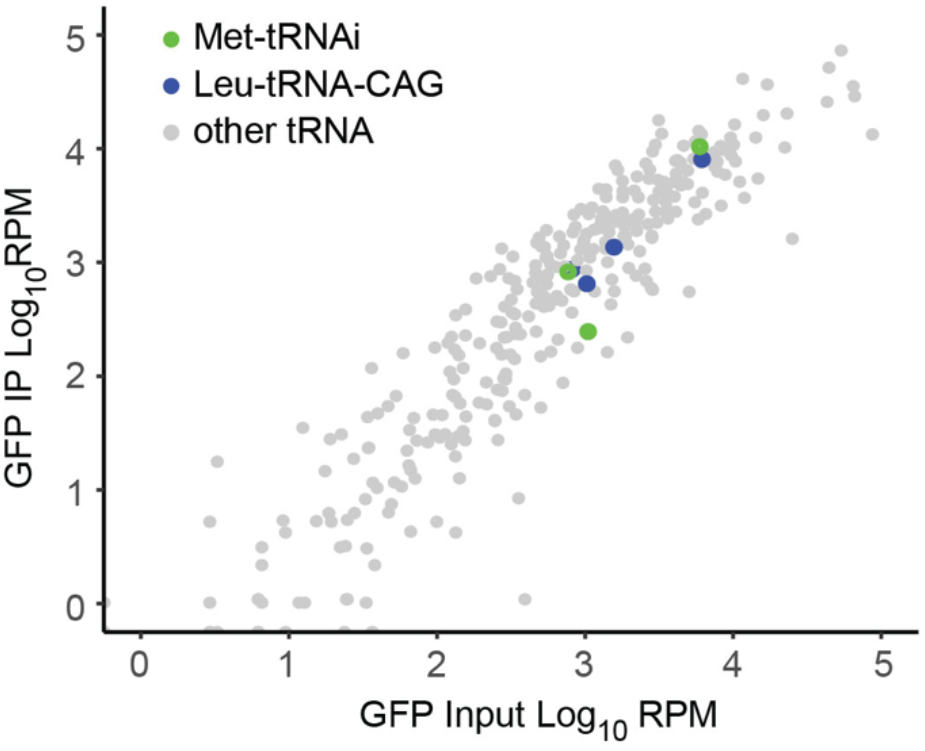
tRNA read counts of GFP input and IP samples. Met-tRNA_i_ and Leu-tRNA-CAG species are indicated in green and blue, respectively.

**Figure S6.**
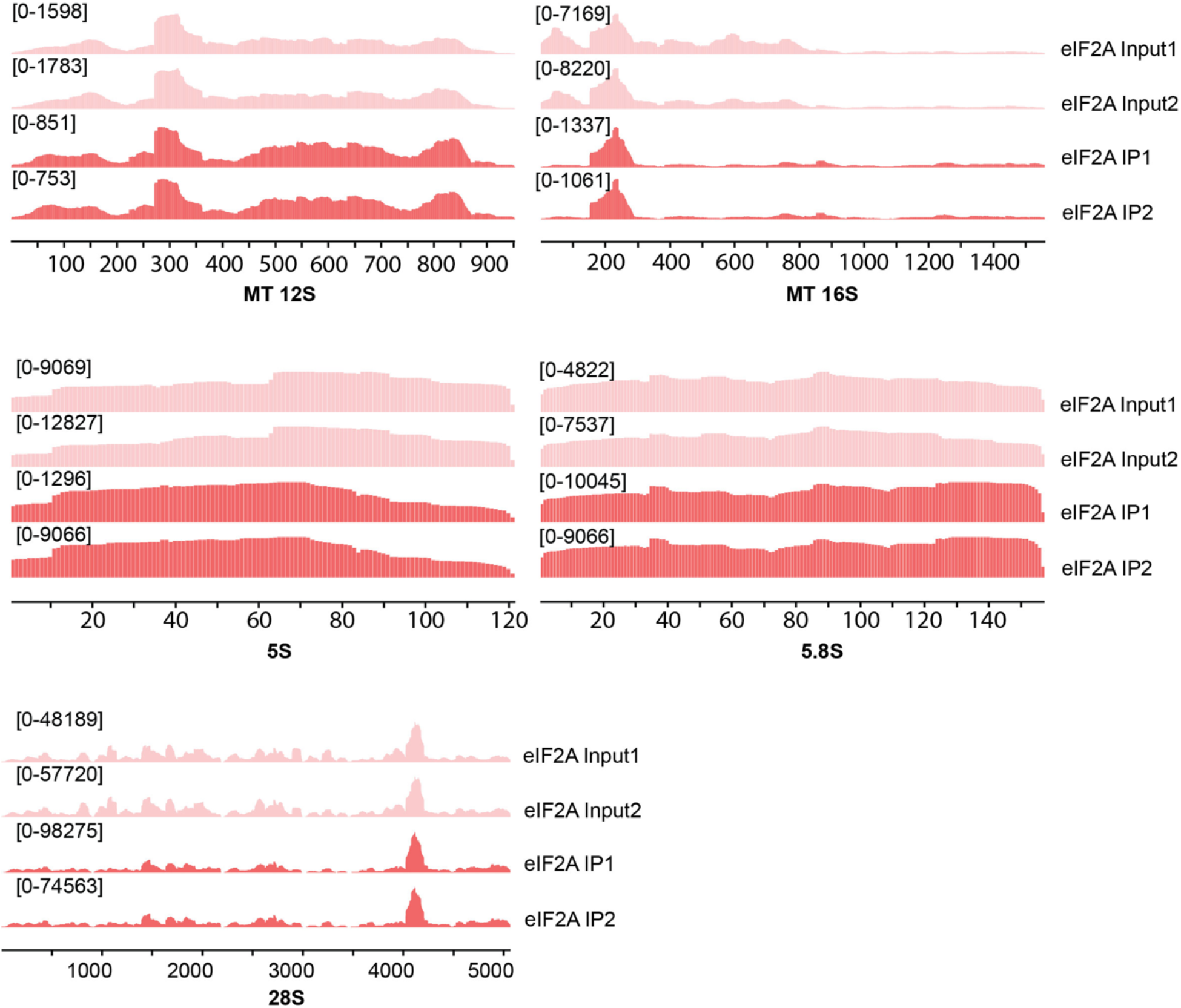
eCLIP read coverage of eIF2A input and IP samples on additional rRNAs.

**Figure S7.**
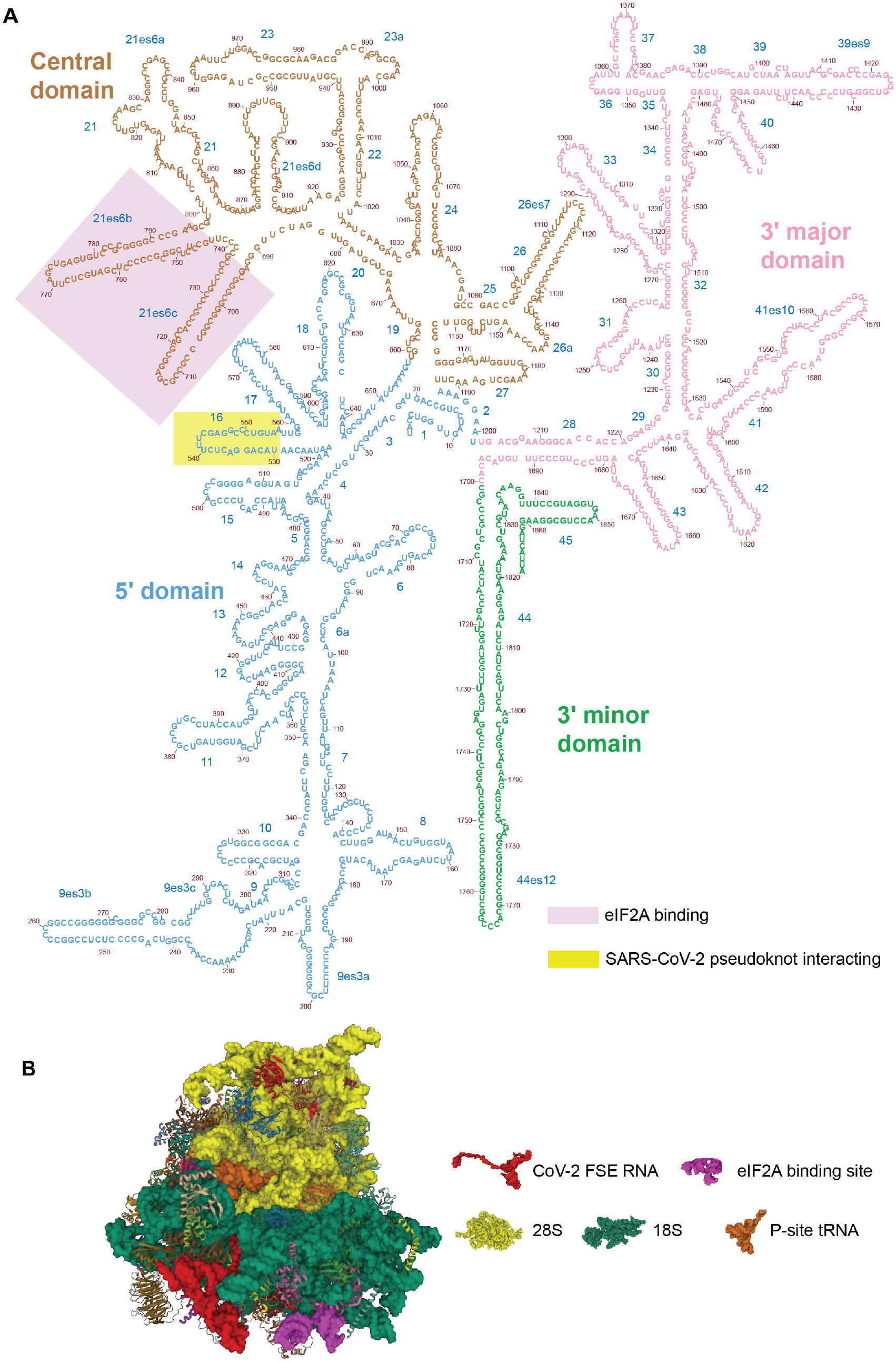
eIF2A binding site on 18S ribosome RNA. (**A**) 2D structure of human 18S ribosome RNA. Interactions with eIF2A and SARS-CoV-2 FSE^3^ are highlighted. Image was modified from RiboVision^39^. (**B**) 3D structure of SARS-CoV-2 RNA and 80S ribosome^3^, with eIF2A binding site indicated in purple.

## Notes

### Competing Interest Statement

The authors have declared no competing interest.

